# The Superior Frontal Longitudinal Tract: Connection between the Dorsal Premotor and the Dorsolateral Prefrontal Cortices

**DOI:** 10.1101/2020.02.01.930552

**Authors:** Mudathir Bakhit, Masazumi Fujii, Ryo Hiruta, Masayuki Yamada, Kenichiro Iwami, Taku Sato, Kiyoshi Saito

**Affiliations:** Department of Neurosurgery, Fukushima Medical University, 1 Hikarigaoka, Fukushima, 960-1295, Japan; Department of Neurosurgery, Aichi Medical University, 1-1 Yazakokarimata, Nagakute, Aichi, 480-1195, Japan

## Abstract

The structural connection between the dorsal premotor (PMd) and the lateral prefrontal cortices (DLPFC) has been revealed, in a few studies, as the frontal longitudinal system (FLS). This study conducted a tractography analysis and a limited, postmortem, white-fiber dissection to investigate the superior FLS tract (SFLT) and analyze both its symmetry and termination point patterns. An analysis of spatial location, termination points, laterality, and correlation with the subjects’ gender or handedness was performed. SFLT was constructed in 100% of right and 88% of left hemispheres. The tracts exhibited variable patterns in different subjects in their posterior terminations. Additionally, the SFLT was found to possess a complex spatial relationship with the adjacent bundles. The SFLT was revealed successfully in two right hemispheres, where the posterior terminations were found to originate in the PMd, and its posterior terminations being totally separate from the superior longitudinal fasciculus.

## Introduction

The brain’s frontal lobe structural connectivity is not perfectly understood due to its complexity, including the connection between the dorsal premotor (PMd) and the dorsolateral prefrontal cortex (DLPFC). Several animal tracing, and functional and probabilistic tractography analysis, reports supported the assumption of its existence (Lu, Preston, & Strick, 1994; Luppino, Rozzi, Calzavara, & Matelli, 2003; Schulz et al., 2019; Tomassini et al., 2007). The term frontal longitudinal system (FLS) was introduced in published studies of the frontal lobe’s white fibers, when it was initially considered a series of U-fibers (Catani et al., 2012; Thiebaut de Schotten, Dell’Acqua, Valabregue, & Catani, 2012). Recently, a fiber dissection study revealed that the U-fibers assumption might be incorrect since the FLS is actually a group of frontal intralobar tracts (Komaitis et al., 2019). Whether the FLS is a separate tract or a continuation of the superior longitudinal fasciculus (SLF) remains debatable. This study hypothesized that the superior bundle of the FLS is a separate, frontal, intralobar tract and not a continuation of the SLF; it is referred to hereafter as the superior frontal longitudinal tract (SFLT). This paper attempts to expand on the works introduced in these recent reports by conducting a thorough virtual dissection of the FLS in a large sample of healthy subjects using generalized, q-sampling imaging (GQI) tractography to study, in detail, its shape, termination patterns and their incidence, and spatial relationship with surrounding structures. It focuses only on the superior or dorsal part: the SFLT (Catani et al., 2012; Komaitis et al., 2019). Furthermore, a white-matter dissection is conducted in order to attempt to confirm its terminations.

## Results

### The dataset

Data of 48 healthy adults from the Human Connectome Project (HCP) database was imported, which included 23 females (47.9%) and 28 right-handed (58.3%) subjects with a mean age of 28.19 ± 3.97 years (range 22–35 years).

### GQI tractography

GQI tractography revealed the SFLT in all 48 (100%) of the right side hemispheres and in 42 (88%) of the left side hemispheres (Fig. 1). The tractography methodology is explained in Fig. 2. Using the parameters presented in the methods section, the streamlines terminated at either the rostral middle frontal (rMFG) or the frontal pole (FP) regions (Fig. 2 B). First, other association, projection, and commissural fibers were removed (Fig. 2 C-F). Then, the remaining bundles were approached from the medial perspective; the select tool in the DSI studio software was used to select the bundle located underneath several superficial U-fibers that was running in a rostro-caudal fashion from the premotor and, sometimes, the motor regions (the precentral gyrus [PCG], the posterior parts of both superior frontal gyrus [SFG], and the caudal middle frontal gyrus [cMFG]) to the DLPFC (Fig. 2 G - red circle). After a few trials, the short U-fibers were eliminated, using the select and remove tools, to construct a separate SFLT (Fig. 2 J). Several cases demonstrated two bundles (one running dorsally and the other ventrally), which were mostly superior and inferior FLS, respectively (Catani et al., 2012; Komaitis et al., 2019). As the inferior FLS is beyond the scope of this study, it was removed.

**Fig. 1.**
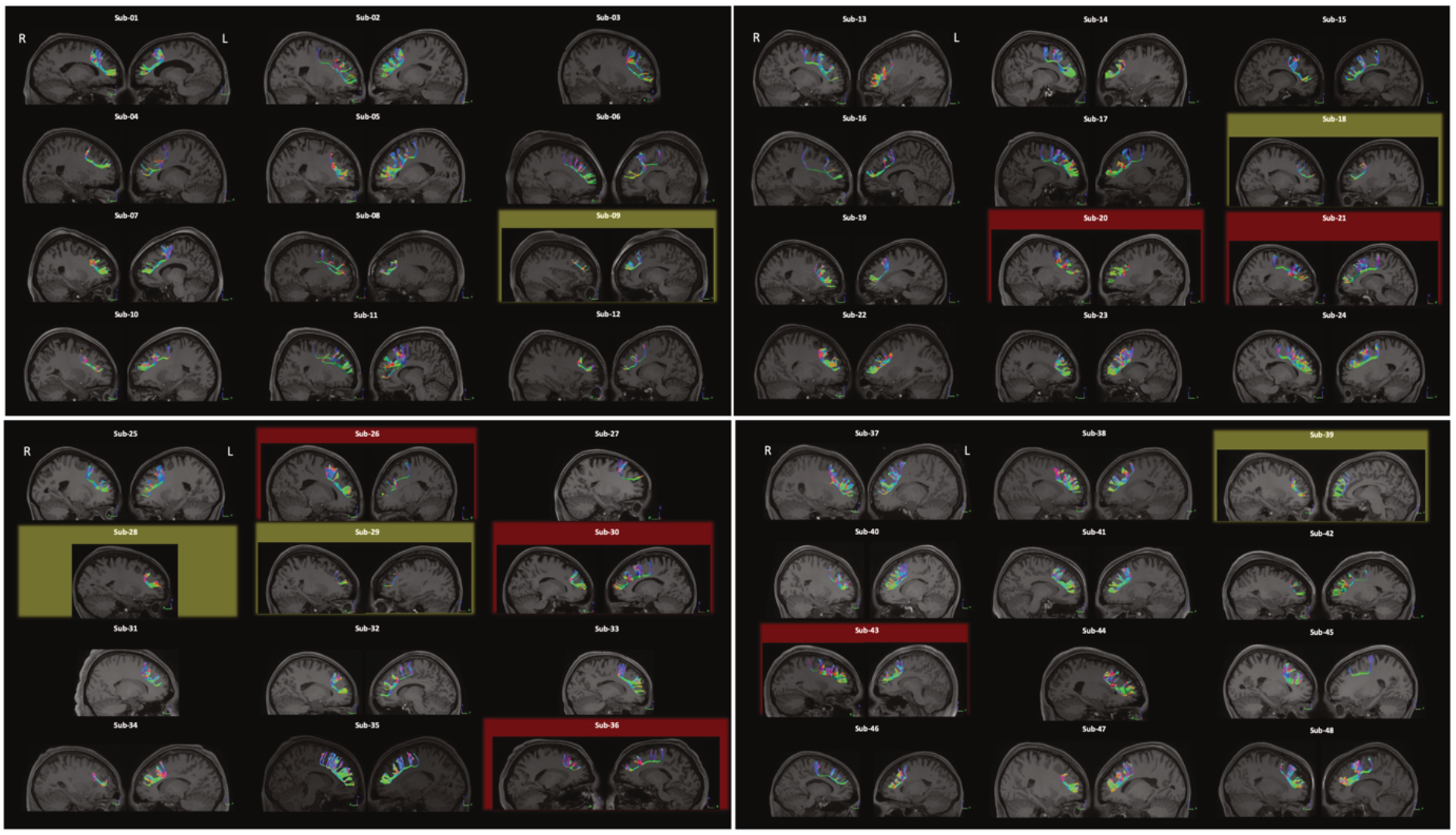
SFLT of subjects 01 to 48. Red highlights represent an existing SFLT made of a chain of two fibers in one side. Yellow highlights represent a lack of posterior termination in the PMd in both sides.

**Fig. 2.**
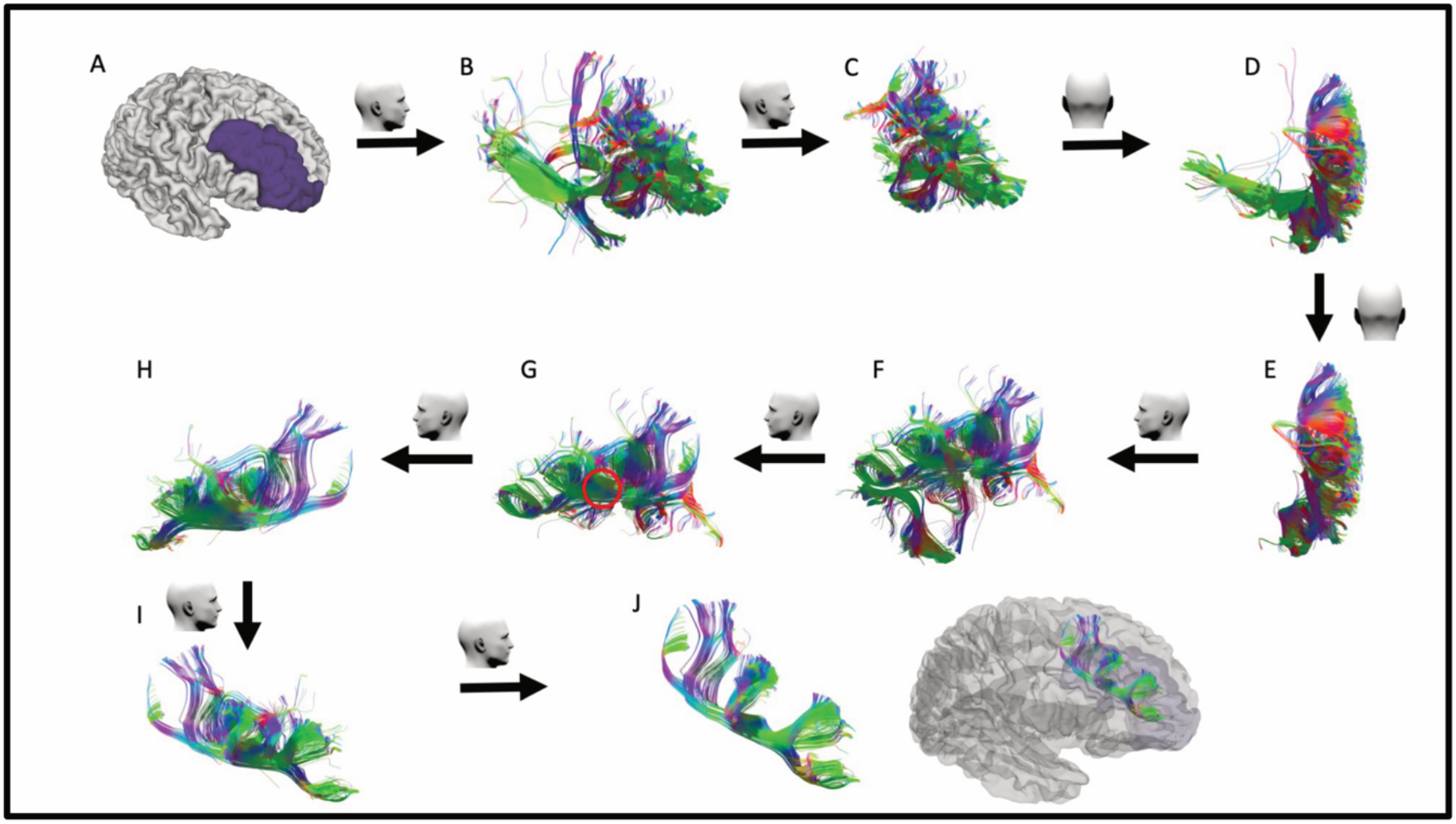
The SFLT construction steps in the right side using DSI studio. (A) The hemispheric white matter ROI are set as a seed region, and the merged rMFG and FP are set as an end region. (B) The raw result of the GQI tractography. (C–F) Irrelevant association, projection, and commissural fibers are removed. (G) The SFLT can be seen from the medial perspective running rostro-caudally (red circle). The select tool was used to focus on this bundle. (H–I) Removal of all remaining U-fibers. (J) A completely constructed SFLT.

### Spatial location

Figure 3 shows the overlap image of the SFLT on both sides in the 48 subjects. The SFLT running path of all subjects lies in the deep white-matter substance of the middle frontal gyrus (MFG) with a maximum frequency at the area antero-lateral to the anterior horn of the lateral ventricles. Figure 3 reveals that the terminations of the overlapping image also exist in the dorsal PCG.

**Fig. 3.**
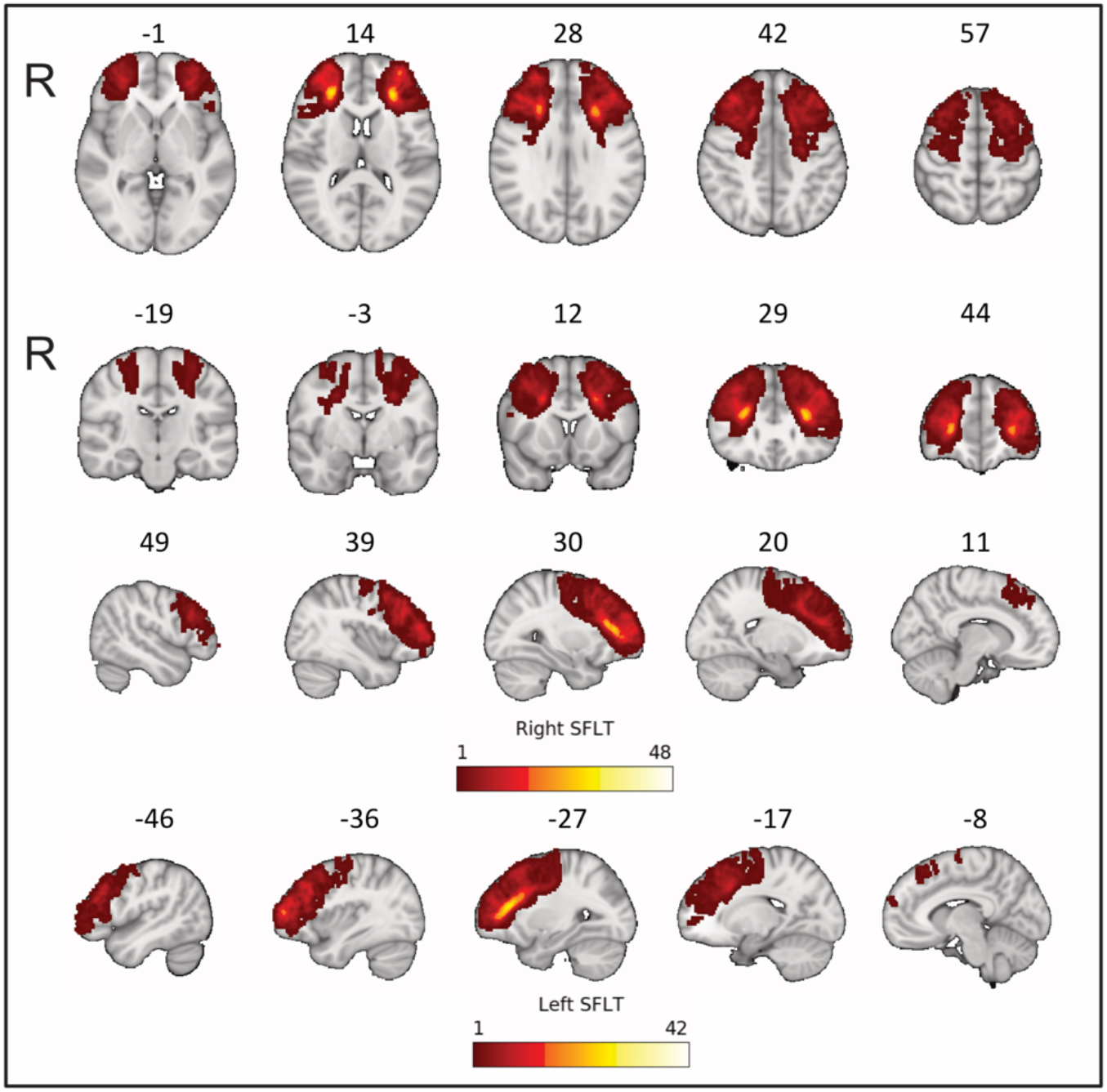
Overlap image of 45 subjects and 48 subjects in the left and right sides, respectively.

### Subcomponents of the SFLT

The SFLT subcomponent regions and probable Brodmann areas (BA) of the terminations are shown in Fig. 4, and an example from one subject is shown in Fig. 5. The GQI tractography of the SFLT revealed variations in the posterior terminations. Conversely, the anterior termination was mostly the same in the rMFG and the FP (BA 9/46/10). The posterior terminations were in the MFG and SFG (BA 6-rostral/8) or the PCG (BA 4/6-caudal). The occurrence rate of these subcomponents varied, as demonstrated in Fig. 4. The MFG subcomponent was present in the left and right sides in 38 (79%) and 45 (94%) subjects, respectively. The PCG subcomponent was present in the left and right sides in 15 (31%) and 11 (23%) subjects, respectively. The SFG subcomponent was present in the left and right sides in 29 (60%) and 21 (44%) subjects, respectively. Five samples revealed no BA6 termination in both sides and were shorter than the other samples (Fig. 1, yellow highlight). Another 12 samples showed unilateral BA6 termination.

**Fig. 4.**
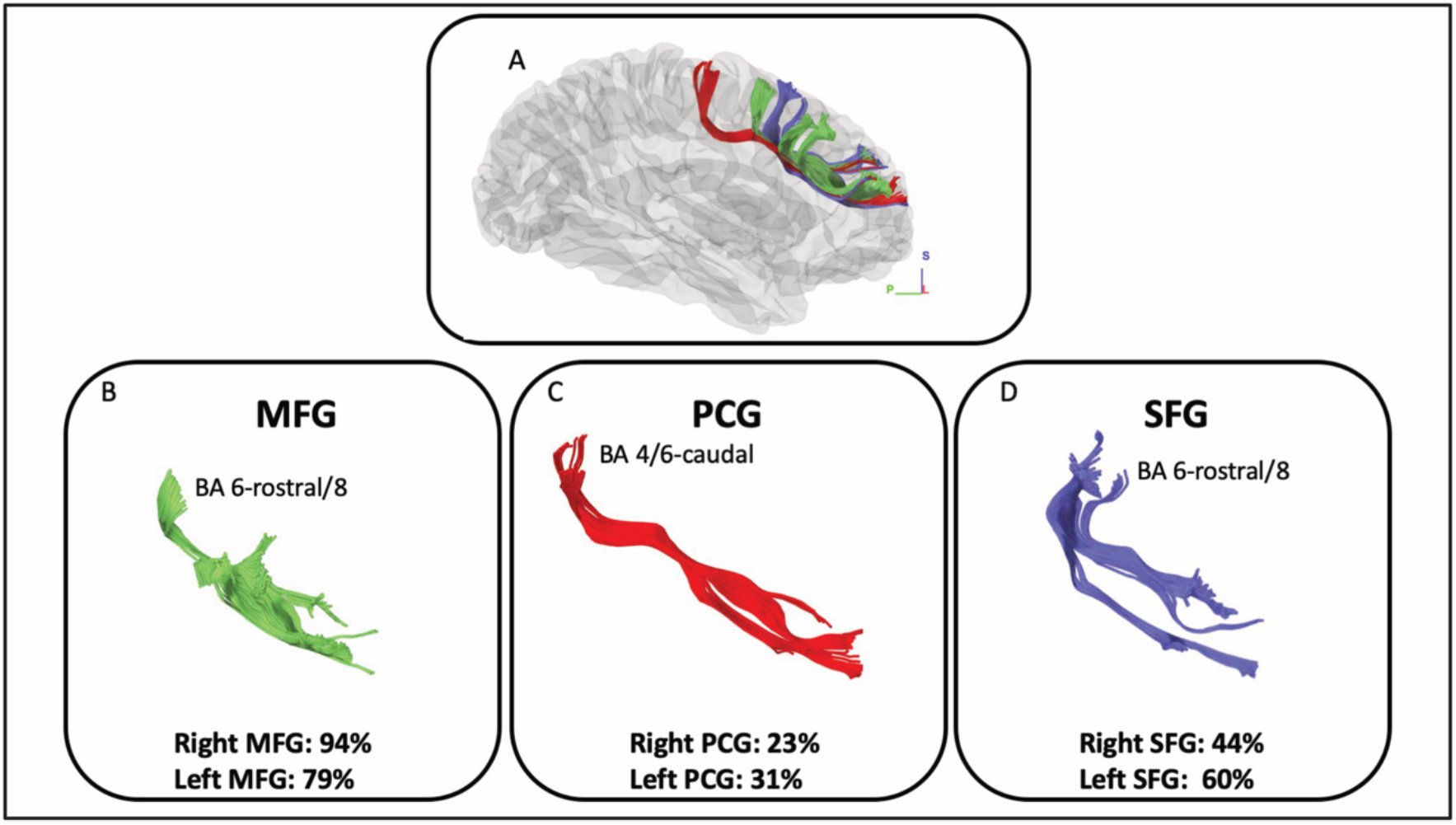
Constructed subcomponents of a right side SFLT in one subject based on posterior termination points as a model example. (A) All subcomponents of the SFLT. (B, C, D) The BA of the termination points and the occurrence rates of the MFG, PCG, and SFG subcomponents, respectively.

**Fig. 5.**
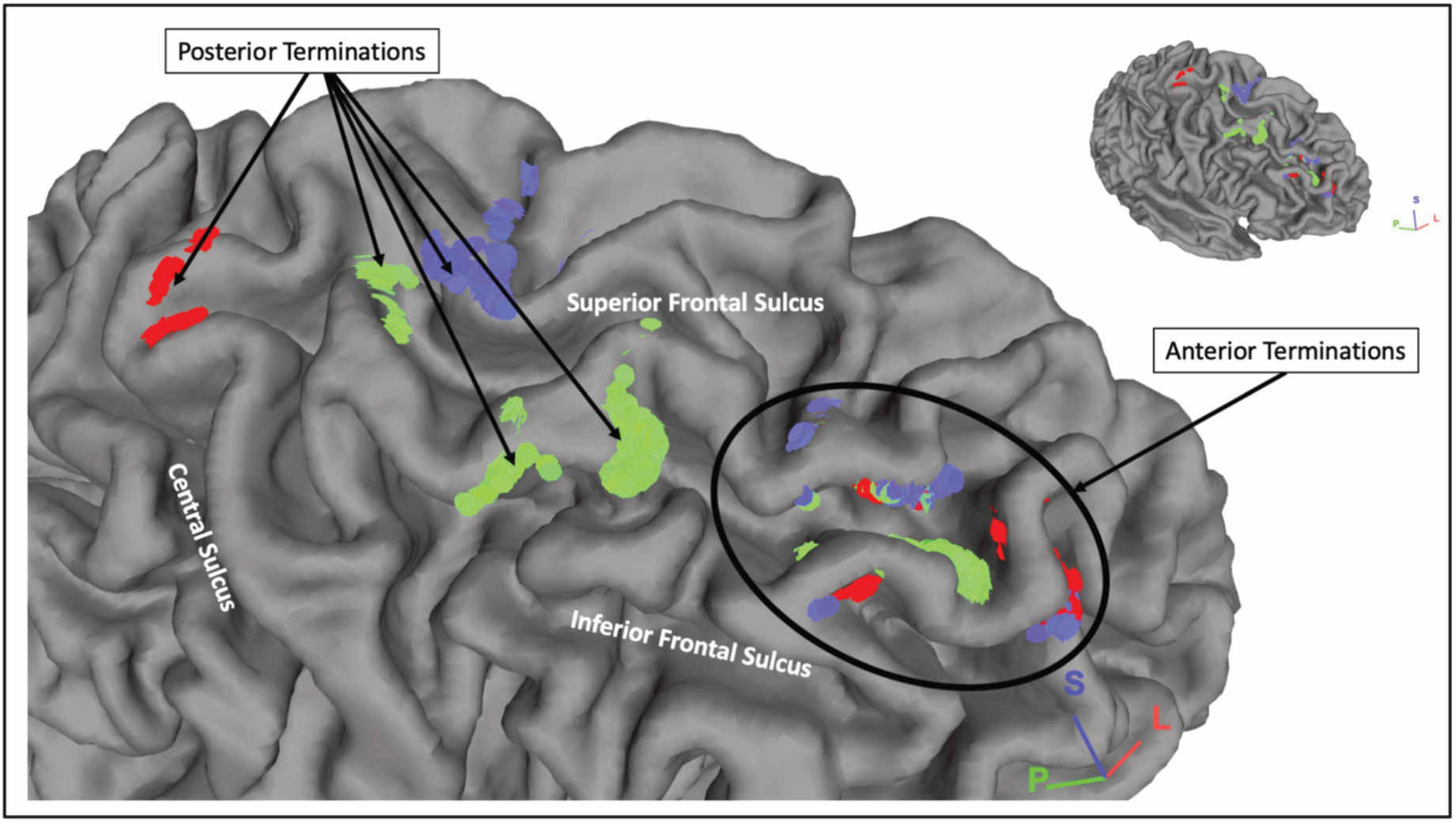
The termination points of a right side SFLT in one subject. The MFG, the PCG, and the SFG subcomponents are shown in green, red, and blue, respectively.

### Spatial relationships with adjacent white matter bundles

A template averaged from a total of 1021 subjects from the HCP was used to study the spatial relation of the SFLT with adjacent white matter bundles as described in Figs. 6 and 7. The SFLT revealed a complex relationship with several adjacently running fibers (association, projection, and commissural). In the left side, the SFLT is closely spatially related to the following tracts: 1) The frontal aslant tract (FAT), which combines with the posterior termination of the SFLT at the PMd area (Fig. 6 E), 2) The arcuate fasciculus, of which the rostral end lies ventral to the PMd segment of the SFLT (Fig. 6 F), 3) The inferior front-occipital fasciculus (IFOF), of which the frontal fibers lie medially to the SFLT (Fig. 6 G), 4) The anterior projection fibers (APF), which lie medially and also share the same plane with the IFOF (Fig. 6 H), 4) The corpus callosum (CC), which intermingles with whole SFLT fibers (Fig. 6 I), and 5) The corticospinal tract (CST), which is a special case only in the left SFLT, where it combines with the superior FLT termination at the dorsal PCG (Fig. 6 J).

**Fig. 6.**
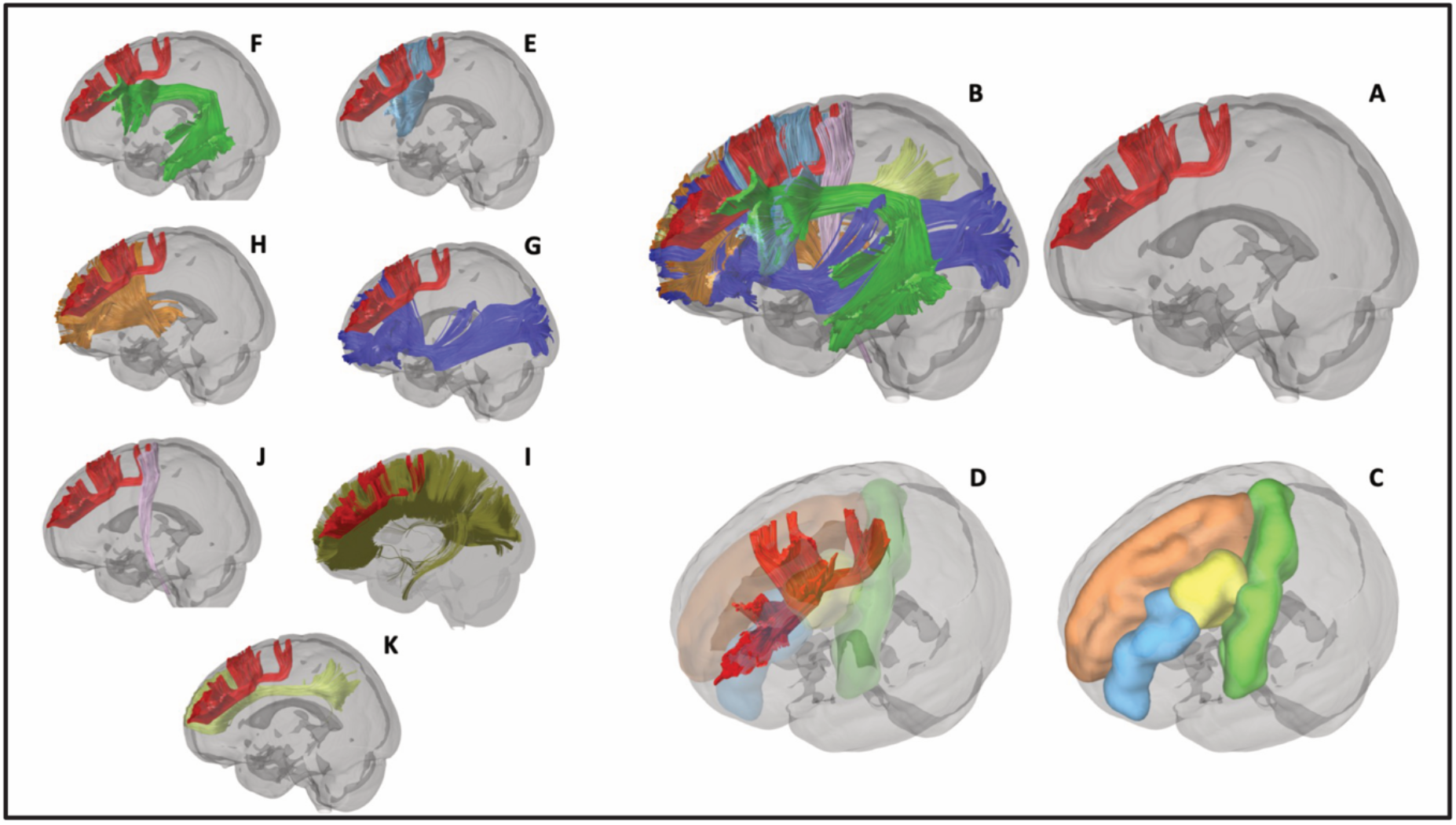
Spatial anatomical relationships of the left SFLT. (A) The constructed SFLT from the HCP-1021 template. (B) The SFLT with all related white-matter fibers (the commissural fibers are not included for the sake of clarity). (C–D) The left SFLT in relation to the PCG (green), SFG (orange), cMFG (yellow), and rMFG (blue). (E) FAT. (F) Arcuate fasciculus. (G) IFOF. (H) APF. (I) Corpus callosum. (J) CST. (K) CB.

**Fig.7.**
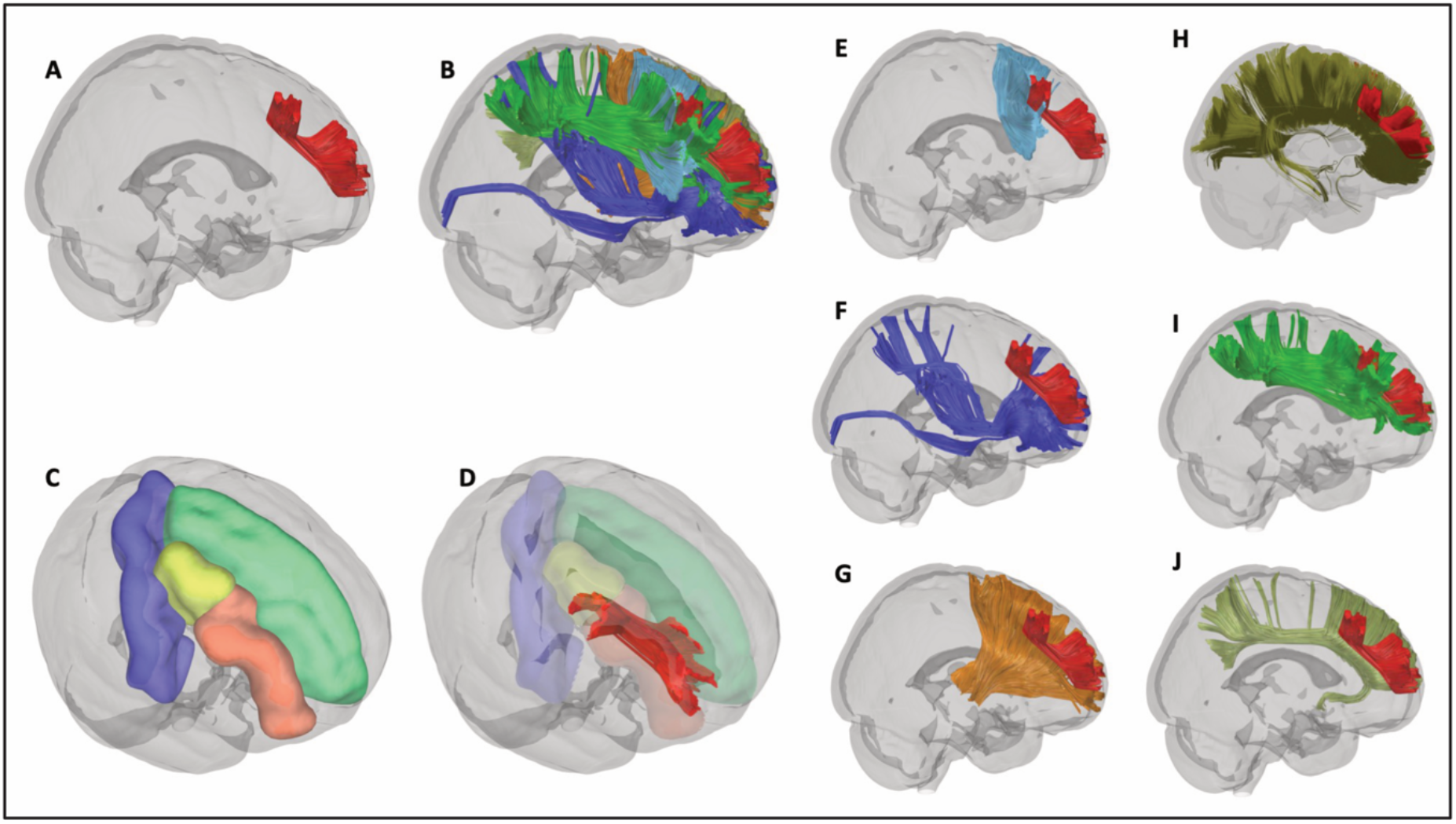
Spatial anatomical relationships of the right SFLT. (A) The constructed SFLT from the HCP-1021 template. (B) The SFLT with all related white-matter fibers (the commissural fibers are not included for the sake of clarity). (C–D) The right SFLT in relation to the PCG (blue), SFG (green), cMFG (yellow), and rMFG (orange). (E) FAT. (F) IFOF. (G) APF. (H) Corpus callosum. (I) SLF. (J) Cingulate bundle.

In the right side, the SFLT is closely spatially related to the following tracts: 1) The FAT, which lies medially and combines with the posterior termination of the SFLT (Fig. 7 E), 2) The IFOF, which has the same location as the left side and lies medially to the SFLT (Fig. 7 F), 3) The APF, which also shares the same position as the left side and lie medially to the SFLT (Fig. 7 G), 4) The CC, which intermingles with whole SFLT fibers (Fig. 7 H), and 5) The SLF, which is a special case in the right SFLT (Fig. 7 I). The SFLT lies within the rostral SLF extension, especially the SLF II. This study also constructed the cingulate bundle (CB) in both sides to distinguish the SFLT from the CB; in both the template and in all subjects, the SFLT lies laterally to the plane of the cingulate gyrus with both the IFOF and the APF between them (Figs. 6 K and 7 J).

In addition, there are some differences between the right and left SFLT based on the HCP-1021 template. The left SFLT has a part that terminates at the dorsal PCG (BA 4/6-caudal) as well as a part that comes from the junction between the SFG and the cMFG (BA 6-rostral/8). The right side has a single bundle with termination posteriorly at the junction between the cMFG and the rMFG (BA 8). Both sides share a similar anterior termination in both sides in the anterior part of the rMFG and the FP (BA 9/46/10).

Furthermore, an example from four subjects is shown in figure 8, where the SFLT is clearly demonstrated as an independent tract and not a continuation of the SLF.

**Fig. 8.**
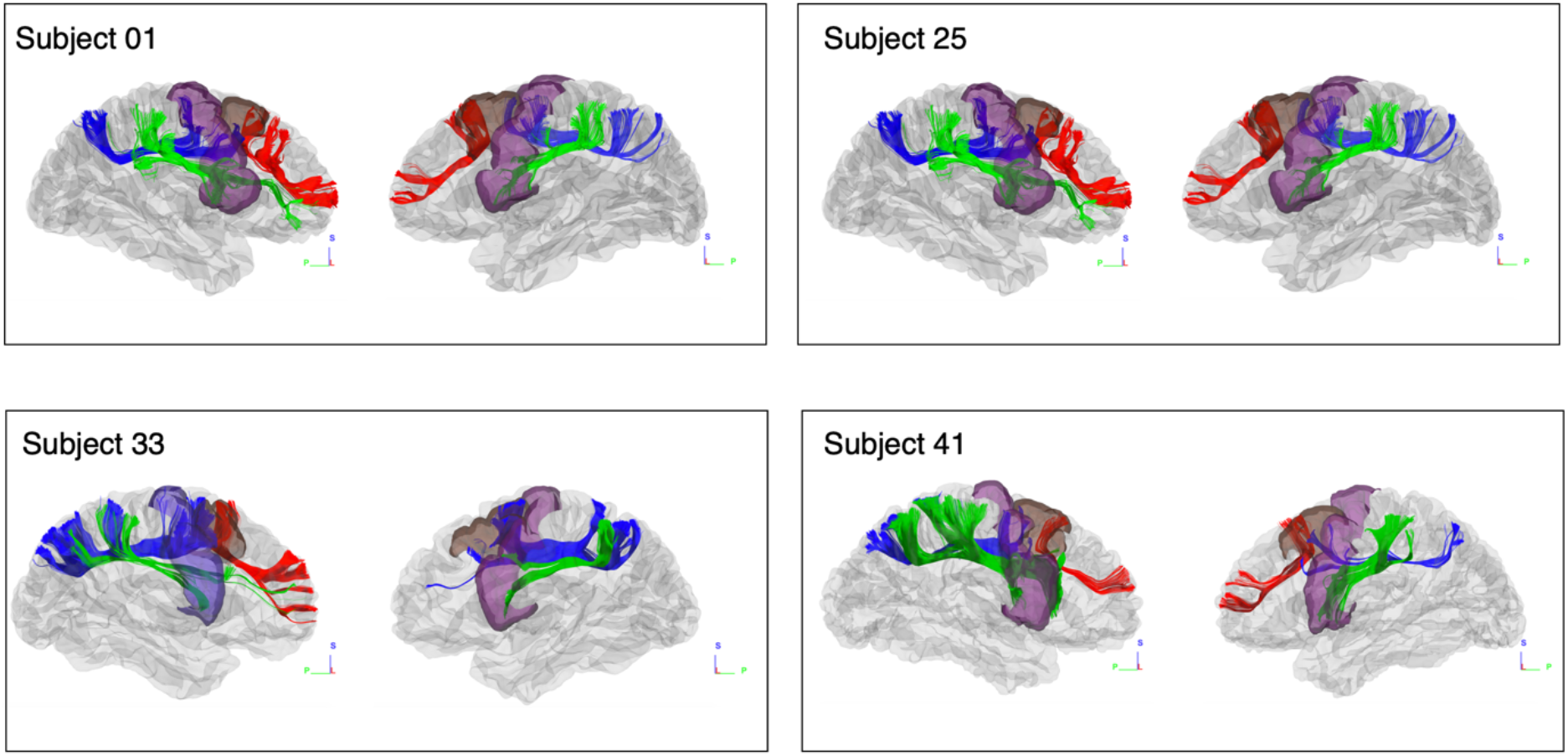
GQI tractography of four subjects, showing the spatial relation between the SFLT (red), SLF 2 (blue), and SLF 3 (green). SFLT were revealed as a separate tract and not a continuation of SLF. The ROIs are: cMFG (brown) and PCG (purple). Subject 33 has only a right side SFLT.

### Statistical analyses of the GQI indices and the laterality index (LI)

The mean values of the SFLT and its subcomponent volume ratios and fraction anisotropy (FA) means are shown in Table 1. The mean value of the whole volume ratio of the SFLT streamlines in the right side and in the left side in the whole number of subjects was 0.66% ± 0.39% and 0.63% ± 0.4%, respectively. Regarding the volume ratio’s LI, 14 (29%) had a right laterality, 12 (25%) revealed a left laterality, and 22 (46%) had undetermined laterality.

**Table 1.** SFLT and its subcomponents: a descriptive analysis

| | Subjects (%) | Mean ± SD | Maximum |
| --- | --- | --- | --- |
| Left SFLT | 42 (88%) |  |  |
| Volume ratio % | | 0.63 ± 0.42 | 1.36 |
| FA mean | | 0.55 ± 0.22 | 0.71 |
| Right SFLT | 48 (100%) |  |  |
| Volume ratio % | | 0.66 ± 0.39 | 1.74 |
| FA mean | | 0.62 ± 0.07 | 0.71 |
| Left MFG subcomponent | 38 (79%) |  |  |
| Volume ratio % | | 0.38 ± 0.31 | 1.03 |
| FA mean | | 0.50 ± 0.27 | 0.71 |
| Right MFG subcomponent | 45 (94%) |  |  |
| Volume ratio % | | 0.50 ± 0.39 | 1.74 |
| FA mean | | 0.58 ± 0.17 | 0.71 |
| Left PCG subcomponent | 15 (31%) |  |  |
| Volume ratio % | | 0.07 ± 0.14 | 0.73 |
| FA mean | | 0.21 ± 0.31 | 0.72 |
| Right PCG subcomponent | 11 (23%) |  |  |
| Volume ratio % | | 0.05 ± 0.11 | 0.5 |
| FA mean | | 0.15 ± 0.27 | 0.7 |
| Left SFG subcomponent | 29 (60%) |  |  |
| Volume ratio % | | 0.22 ± 0.25 | 0.95 |
| FA mean | | 0.37 ± 0.31 | 0.69 |
| Right SFG subcomponent | 21 (44%) |  |  |
| Volume ratio % | | 0.15 ± 0.19 | 0.49 |
| FA mean | | 0.25 ± 0.31 | 0.66 |

The mean value of the whole SFLT streamlines’ FA mean in the right side and in the left side in the whole number of subjects was 0.62% ± 0.07% and 0.55% ± 0.22%, respectively. Regarding the FA mean’s LI, 6 (13%) had a right laterality and 42 (88%) revealed undetermined laterality. No subject had a left FA mean laterality. Neither the SFLT nor its subcomponent volume ratio’s LI or its FA mean’s LI correlated with the subjects’ gender or handedness groups (p > 0.05) (Table 2).

**Table 2.** LI pattern distribution and correlation with gender and handedness.

| Tract | LI Pattern | Count | Gender |  |  | Handedness |  |  |
| --- | --- | --- | --- | --- | --- | --- | --- | --- |
|  |  | Total (%) | Female | Male | p | Right | Left | p |
| SFLT |  |  |  |  |  |  |  |  |
| Volume ratio LI | Left | 12 (25%) | 6 | 6 | 0.53* | 7 | 5 | 0.40* |
|  | Undetermined | 22 (46%) | 12 | 10 |  | 8 | 14 |  |
|  | Right | 14 (29%) | 5 | 9 |  | 5 | 9 |  |
| FA mean LI | Left | - | - | - | 0.67† | - | - | 1.0† |
|  | Undetermined | 42 (88%) | 21 | 21 |  | 18 | 24 |  |
|  | Right | 6 (13%) | 2 | 4 |  | 2 | 4 |  |
| MFG subcomponent |  |  |  |  |  |  |  |  |
| Volume ratio LI | Left | 10 (21%) | 5 | 5 | 0.99* | 5 | 5 | 0.58* |
|  | Undetermined | 17 (35%) | 8 | 9 |  | 8 | 9 |  |
|  | Right | 21 (44%) | 10 | 11 |  | 7 | 14 |  |
| FA mean LI | Left | 3 (6%) | 1 | 2 | 1.0† | 1 | 2 | 1.0† |
|  | Undetermined | 35 (73%) | 17 | 18 |  | 15 | 20 |  |
|  | Right | 10 (21%) | 5 | 5 |  | 4 | 6 |  |
| PCG subcomponent |  |  |  |  |  |  |  |  |
| Volume ratio LI | Left | 13 (27%) | 8 | 5 | 0.32* | 6 | 7 | 0.92* |
|  | Undetermined | 25 (52%) | 12 | 13 |  | 10 | 15 |  |
|  | Right | 10 (21%) | 3 | 7 |  | 4 | 6 |  |
| FA mean LI | Left | 12 (25%) | 7 | 5 | 0.73† | 6 | 6 | 0.66† |
|  | Undetermined | 28 (58%) | 13 | 15 |  | 10 | 18 |  |
|  | Right | 8 (17%) | 3 | 5 |  | 4 | 4 |  |
| SFG subcomponent |  |  |  |  |  |  |  |  |
| Volume ratio LIz | Left | 19 (40%) | 8 | 11 | 0.79† | 8 | 11 | 0.80† |
|  | Undetermined | 22 (46%) | 11 | 11 |  | 10 | 12 |  |
|  | Right | 7 (15%) | 4 | 3 |  | 2 | 5 |  |
| FA mean LI | Left | 15 (31%) | 7 | 8 | 1.0† | 5 | 10 | 0.64† |
|  | Undetermined | 27 (56%) | 13 | 14 |  | 13 | 14 |  |
|  | Right | 6 (13%) | 3 | 3 |  | 2 | 4 |  |

\*Chi-squared test, †Fisher's exact probability test

To classify the SFLT into types, the MFG and SFG subcomponents were merged as a single variable since they were likely to share the same BA (Table 3). A two-step cluster analysis was conducted, based on the presence or absence of a subcomponent with a posterior termination at the BA 4/6-caudal (PCG subcomponent) and the BA 6-rostral/8 (MFG and SFG subcomponents). The outcome was four types of SFLT (as presented in Figure 9: Silhouette measure = 0.9). The SFLT types did not exhibit a correlation with the subjects’ gender or handedness groups (p > 0.05) (Table 4).

**Table 3.** SFLT BA subcomponent frequency.

|  | Current study | Komaitis et al. (2019) |
| --- | --- | --- |
| Left BA 4/6-caudal | 15 (31%) subjects | 6 (86%) left hemispheres |
| Left BA 6-rostral/8 | 41 (85%) subjects | 7 (100%) left hemispheres |
| Right BA 4/6-caudal | 11 (23%) subjects | 4 (50%) right hemispheres |
| Right BA 6-rostral/8 | 48 (100%) subjects | 8 (100%) right hemispheres |

**Table 4.** SFLT type distribution and correlation with gender and handedness groups.

| SFLT Type | Count | Gender |  |  | Handedness |  |  |
| --- | --- | --- | --- | --- | --- | --- | --- |
|  | Total (%) | Female | Male | p | Right | Left | p |
| Type 1 | 19 (39.6%) | 10 | 9 | 0.86<sup>†</sup> | 8 | 11 | 1.0<sup>†</sup> |
| Type 2 | 11 (22.9%) | 4 | 7 |  | 4 | 7 |  |
| Type 3 | 7 (12.5%) | 3 | 4 |  | 3 | 4 |  |
| Type 4 | 11 (22.9%) | 6 | 5 |  | 5 | 6 |  |

<sup>†</sup>Fisher's exact probability test

**Fig. 9.**
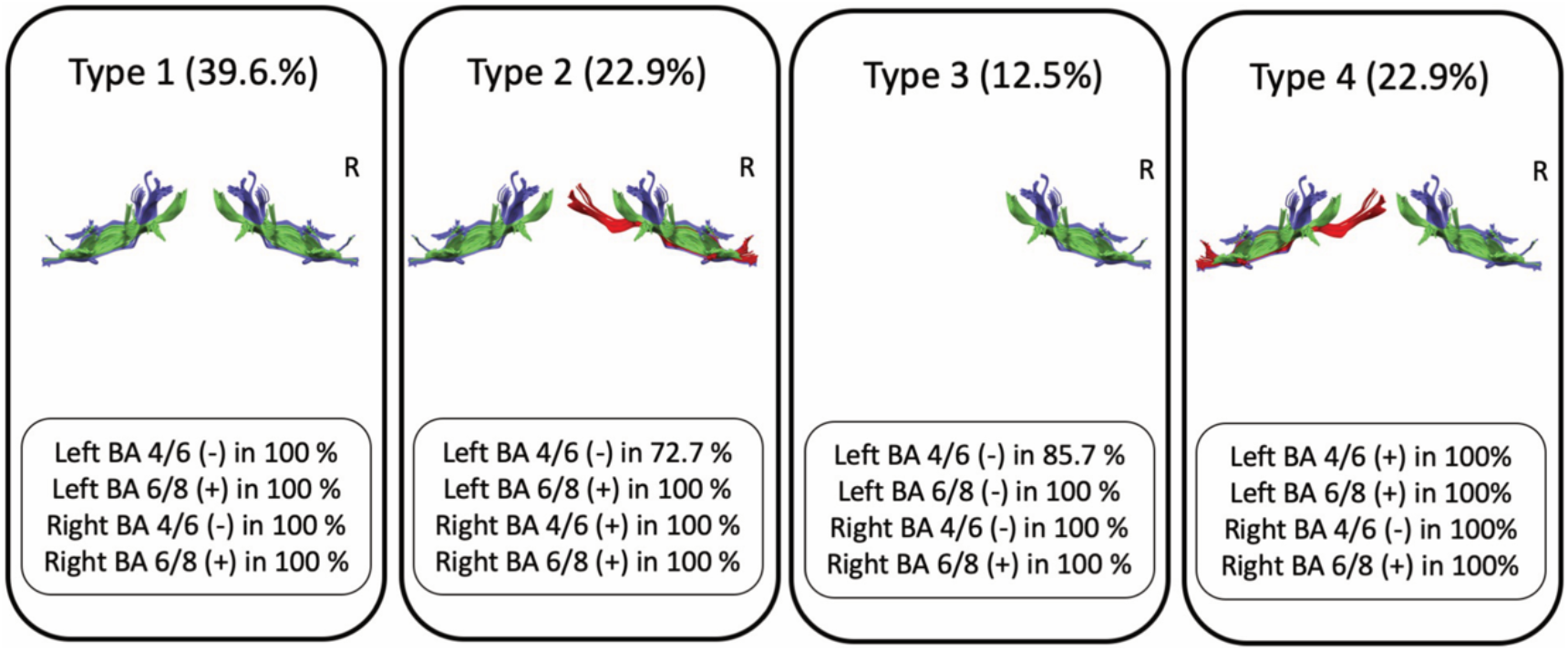
Types of SFLT. The SFLT was divided based on its posterior termination as either “motor segment” arising from the PCG (color) or “premotor segment” arising from the MFG and/or SFG (color). (+) present, (−) absent, BA Brodmann area.

### White-fiber dissection

Based on the prior knowledge of the results obtained using tractography, the dissection was planned first, to start from a point posterior to the fronto-marginal sulcus and second, to dissect in the direction towards the cMFG (Fig. 10 A). After carefully removing the U-fibers in the two right samples, the dissection followed the tracts running rostrocaudally beneath the MFG substances and toward the termination area, which was found to be at the cMFG in the first sample (BA 6-rostral) (Fig. 10 C). Additionally, in this sample, the examination explored the anatomical relationship with the adjacently running SLF and confirmed both that each bundle possessed its own separate posterior terminations and that the SFLT in this sample was not a continuation of the SLF (Figs. 10 D, E). In the second sample, the dissection was extended posteriorly to reach the PCG. A thin band of white fibers was observed to run from the cortex of the transitional zone between the PCG and the cMFG (BA 6-caudal), which curves anteriorly beneath the MFG substances towards the DLPFC and the FP (Figs. 11 A-C).

**Fig 10.**
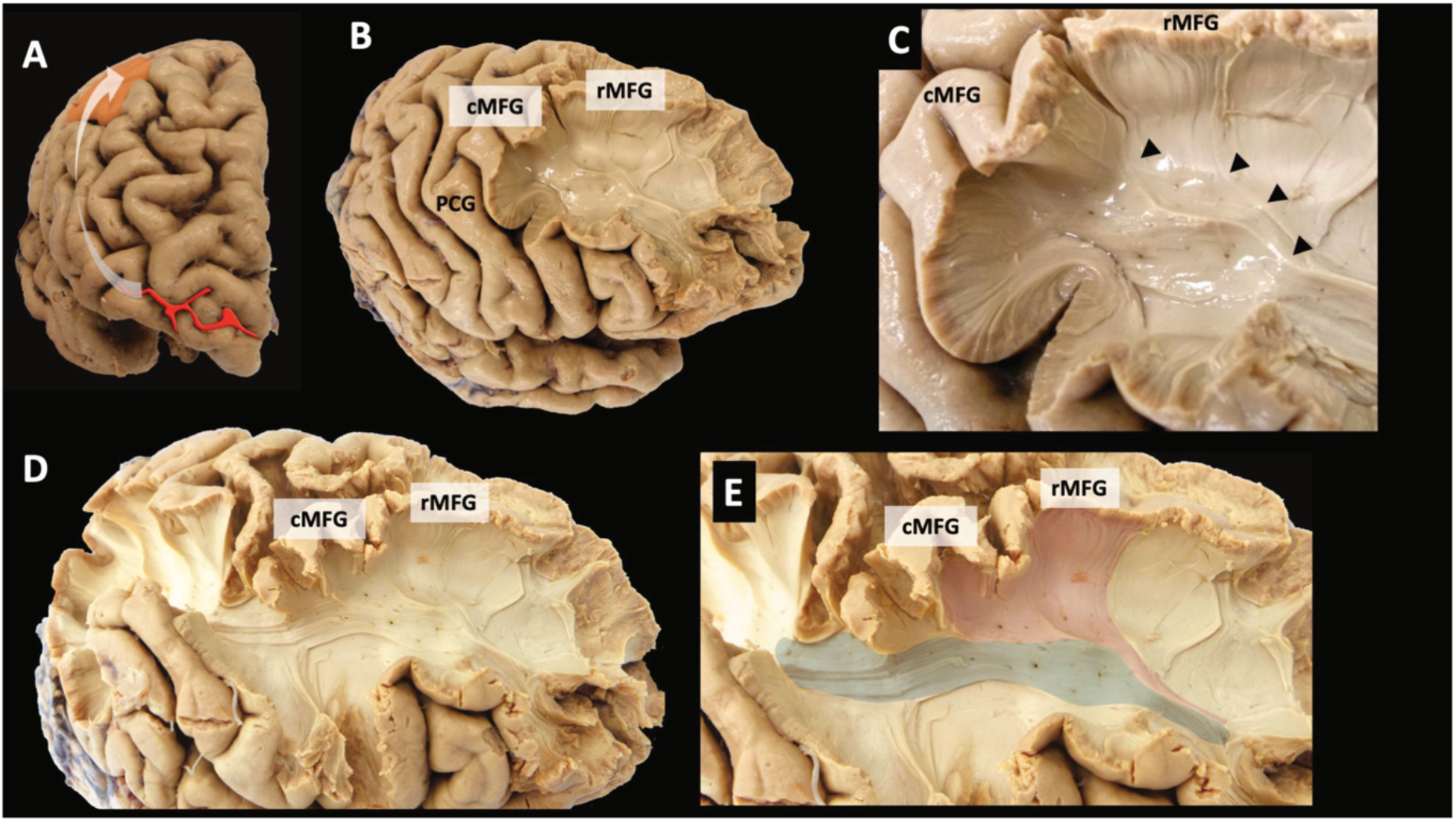
White-matter dissection of a right hemisphere (sample 1). (A) The dissection line starts posterior to the fronto-marginal sulcus (red line) towards the cMFG (orange color). (B) Removing the U-fibers at the MFG reveals the SFLT running beneath. (C) A closer look at the posterior end of the SFLT and how it arises from the MFG cortical region. (D–E) The spatial relationship between the SFLT and SLF is demonstrated.

**Fig. 11.**
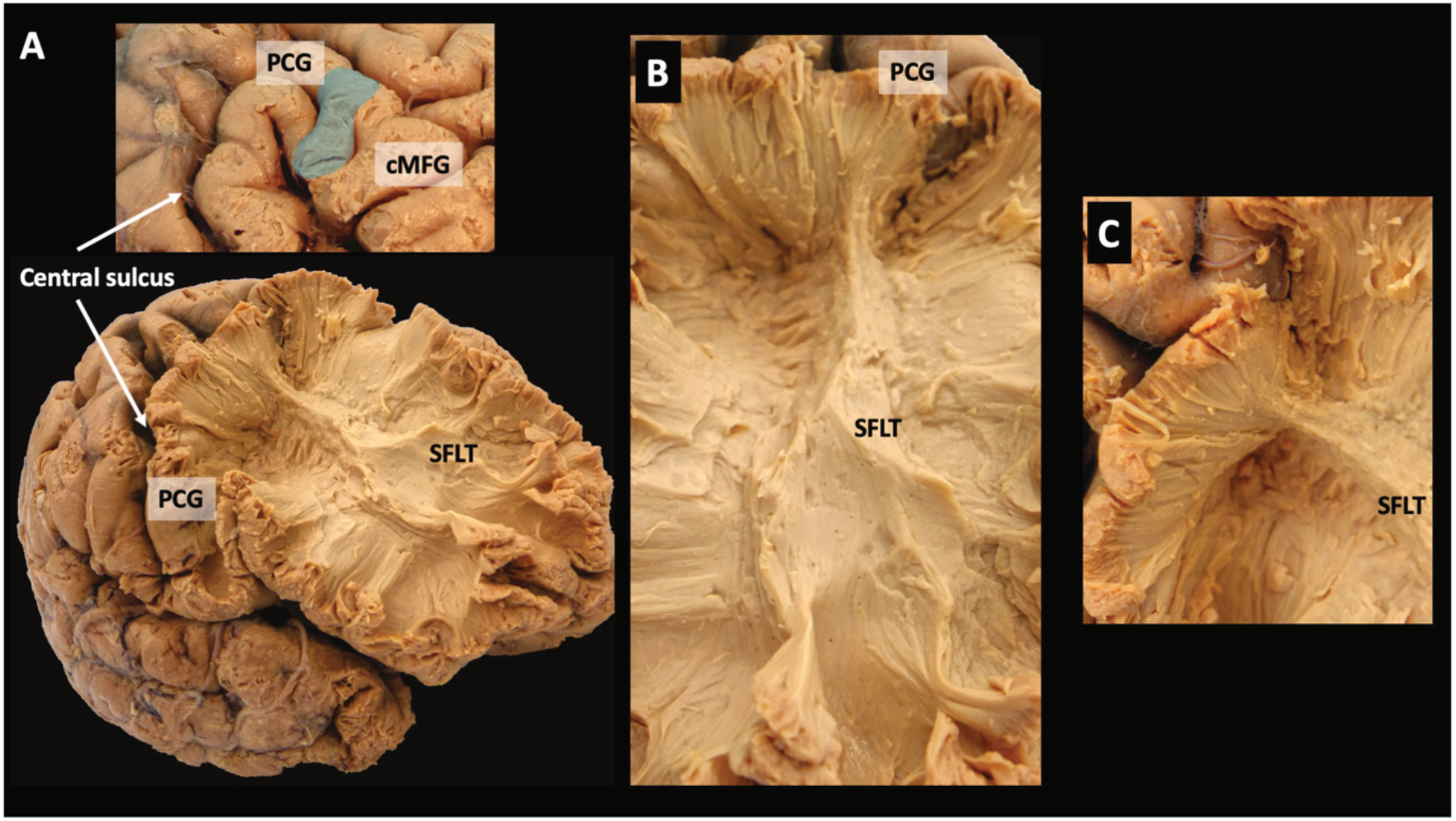
White-matter dissection of a right hemisphere (sample 2). (A) After dissection of most of the frontal lobe and sparing a limited part of the PCG, the SFLT posterior end is revealed, as it arises from the transition zone between the PCG and the cMFG (blue shade). (B) A closer image showing the entire SFLT from an anterior angle. (C) A closer image at the posterior end of the SFLT.

## Discussion

This paper extensively studied the virtual anatomy of the SFLT using GQI tractography with a limited white-fiber dissection, which focused on the anatomical terminations and symmetry; this aspect received little attention in previous reports (Catani et al., 2012; Komaitis et al., 2019; Schulz et al., 2019). In all the samples studied, the SFLT was found to be a frontal intralobar tract, which differs to some previous reports. In their analysis of the frontal short fibers, Catani et al. described the FLS as a chain of U-shaped connections that resemble a prolongation of the SLF connecting the premotor cortex to the DLPFC (Catani et al., 2012). In this study’s opinion, the advantage of processing a multi-shell diffusion image with a higher directional sample (more than 90) and the ability to use modern tractography models (such as GQI) facilitated a better outcome in the tract-rendering process and overcame several potential challenges (such as the effects of crossing and kissing fibers).

The status of the SFLT or the FLS as more than mere U-fibers was confirmed by the extensive microdissection study, recently presented by Komaitis et al. (Komaitis et al., 2019). They recorded the FLS in all 15 brain samples that were studied, which represents a similar proportion (100% and 88% of the right and left hemispheres, respectively) to that achieved in the present study using tractography (Fig. 1).

Furthermore, Komaitis et al. found that, in 80% of samples where a prominent middle frontal sulcus was present, the FLS was encountered just under the most superficial U-fibers at the depth of the sulcus. In the remaining 20% of cases, where the middle frontal sulcus was absent, the FLS was seen to lie just medially to the MFG’s superficial U-fibers. In addition, the FLS fibers were observed to travel at two discrete levels; the superior and the inferior FLS, as described by Catani et al (Catani et al., 2012).These descriptions of the relationship of the FLS to the superficial U-fibers correspond with that observed during this study’s tract construction process (Fig. 2). Furthermore, the location of the SFLT was studied in 48 subjects, and the site was mapped in an MNI space template (Fig. 3). This study is in agreement with Komaitis et al.; it is considered that the SFLT’s maximum intensity lies beneath the MFG’s substances and is located antero-laterally to the anterior horn of the lateral ventricles. The constructed tracts shared mainly the same location of the SFLT body, but they varied in their terminations, especially the posterior part.

The SFLT termination points were studied in the subjects’ native space using the FreeSurfer Desikan-Killiany (DK) atlas as a reference. In agreement with both the spatial location analysis above and the results from the Komaitis et al. study, the present study’s data indicates a similar anterior termination point (in the rMFG at BA 9/46). There was a significant variation between subjects regarding the posterior termination points, which were located in one of three regions: the MFG, the PCG, and the SFG (Figs. 4 and 5). Therefore, in regard to incidence, it was revealed that the MFG is the most frequently occurring subcomponent among the three with a right side preference, and the PCG and the SFG have left side preferences even though the PCG subcomponent has far less incidence. Although this study classified these subcomponents according to their termination in the DK atlas, most subjects demonstrated that the MFG and the SFG subcomponents were significantly adjacent within one probable zone of the BA (the BA 6-rostral/8), while the PCG subcomponent was separate and (when available) situated in the BA 4/6-caudal. The BA arrangement was chosen in this study as a variable in the subsequent cluster analysis.

This paper reviewed an already published HCP-1021 average tractography template (F.-C. Yeh et al., 2018), which revealed differences in shape between the right and left sides; the left side’s SFLT was the only one with both the PCG and SFG subcomponents, while the right side’s SFLT was mainly an exclusive MFG subcomponent. This corresponds to the incidence rate findings since the PCG and SFG subcomponents revealed a higher incidence in the left side (Tables 1 and 2). Moreover, the template explored the surrounding white matter, and the SFLT on both sides revealed a complex close relationship with several white-matter bundles (as shown in the results section and in Figures 6 and 7). The major difference between both sides in this aspect was the relationship between the PCG subcomponent and the CST (which was absent for the right side) and the relationship between the right SFLT (the MFG subcomponent) with the SLF (which was absent for the left side) in the template example (Figs. 6 and 7). In 20% of the hemispheres studied by Komaitis et al., the SLF and the FLS were recorded as two completely distinct tracts in a ratio far lower than that discovered by the present study since it was possible to construct the SFLT in 100% of the right hemispheres. Komaitis et al. reported that, in most (80%) cases, they identified the FLS as the anterior extension of the SLF’s fiber system. In contrast to these results, all the SFLTs constructed in this research were located within the frontal lobe and shared no termination pattern like the familiar ones usually found in the SLF (Fig. 1 and 8) (de Schotten et al., 2011; Wang et al., 2016).

The SFLT (as a whole) or its subcomponents demonstrated no specific laterality in either volume ratio or FA mean values, and most subjects possessed an undetermined pattern. One exception was the MFG subcomponents’ volume ratio LI (Table 2). The latter showed 44%, 21%, and 17% of the subjects with a right side, a left side, and an undetermined LI, respectively. A cluster analysis was undertaken to reveal any possible SFLT subtypes based on these subcomponents; after rearranging them according to their probable BA, four different types (shown in Figure 9) and 39.6% of the population were projected to have a bilateral SFLT with a posterior termination from BA 6-rosrtal/8 (Type 1) only, while 12.9% were assumed to possess only a unilateral right side tract with the same termination (Type 3). Types 2 and 4 have a bilateral BA 6-rosrtal/8 and a unilateral BA 4/6-caudal subcomponent in the right and left sides, respectively. No subtype expressed the co-existence of all the BA subcomponents bilaterally.

This study concluded with a limited white-matter dissection. The SFLT was located in both right samples, and their posterior terminations were followed: one at the MFG (BA 6-rostral/8) in the first sample (Fig. 10) and another at a zone between the PCG and the cMFG (BA 6-caudal) (Fig. 11). In the latter, the distinctiveness of the posterior terminations between the SFLT and the adjacently running SLF was confirmed.

Regarding the BA subcomponents, there is limited similarities between this study’s tractography results (incidence in subjects) and the white-fiber dissection study (incidence in hemispheres) undertaken by Komaitis et al. The right BA 6-rostral/8 subcomponent was found in 100% of cases in both studies. The remaining BA distribution did not show other similarities and can be seen in Table 3.

Based on this study’s findings and previously published research, the SLFT represents a connection between the DLPFC and the PMd cortex. The BA4/6-caudal subcomponent seems to exist genuinely in humans at a much lower rate than BA6-rostral/8, which might explain why other similar studies in humans could not confirm its existence (Schulz et al., 2019). Still, we do not have a clear understanding of why the BA4/6-caudal subcomponent has such a low rate in tractography construction, However, such an interpretation should be made with care since no direct connection between the caudal PMd (PMd proper) and the DLPFC was detected in animal studies (Geyer, Matelli, Luppino, & Zilles, 2000; Lu et al., 1994). Also, we should keep in mind that tractography is not a perfect technique, and errors can happen. The available data concerning a human direct structural pathway between the rostral PMd and the DLPFC is still limited, apart from a few functional studies involving healthy participants (Tomassini et al., 2007), well-recovered chronic stroke patients (Schulz et al., 2019), and postmortem examinations (Komaitis et al., 2019). Regarding the motor network, the DLPFC has been reported as primarily connected to the PMd area, the ventral premotor area, and the supplementary motor area. Brain-tracing studies in animals have revealed similar connections (Geyer et al., 2000; Lu et al., 1994; Luppino et al., 2003).

Regarding the SFLT’s possible functional rule, crucial structures for motor and cognitive skills are concomitantly activated during motor and cognitive tasks, including the PMd and the DLPFC (Diamond, 2000; Hanakawa, 2011). The activation of these brain structures is stronger in difficult (as opposed to easy) tasks (Diamond, 2000). When performing a difficult (i.e., complex) motor task, higher-order cognitive resources (such as executive tasks) are required to perform the action successfully. Furthermore, the rostral PMd (pre-PMD) and the DLPFC may contribute to executive function through sequence generation and attentional selection, respectively, and the functional coupling between them seems to play a pivotal part in integrating these executive processes. (Abe et al., 2007). Therefore, the SFLT might serve as the structural backbone in these functions. Additional studies involving functional connectivity and task-related functional MRI (fMRI) analysis regarding the SFLT’s spatial correlation might be necessary to confirm this assumption.

As demonstrated in this work, the superior part of the FLS is neither a system comprising U-fibers nor a continuation of another white-matter bundle. Instead, it is a fully structured white-fiber bundle; consequently, it is proposed that it is identified as the superior frontal longitudinal tract.

Some limitations of the present study are considered below. A few of the constructed SFLTs were significantly shorter and lacked a BA 6 termination but, based on their spatial location, they were identified as SFLT rather than being discarded (images highlighted in yellow in Figure 1). Additionally, six subjects who had a unilateral SFLT comprising a chain of two fibers (images highlighted in red in Figure 1) were also included as the SFLT in this study since they shared the same spatial location. Furthermore, the current work focused only on the SFLT and ignored the inferior part of the FLS, as it was assumed the multiple dissimilarities might require a separate dedicated study. The reason of much lower rate of the BA4/6-caudal subcomponent is not clear for us, and factors related to the tractography cannot be excluded. Finally, the current work focused on the structure of the SLFT and did not include any functional imaging analysis.

In conclusion, this study demonstrated that the SFLT is a frontal intralobar tract connecting the PMd to the DLPFC, not a mere continuation of the SLF. The tract exhibited variable patterns in its posterior terminations among the samples. It revealed a complex and close spatial relationship with several white-matter bundles including, the FAT, the arcuate fasciculus, the IFOF, the CST, the CC, and the SLF.

## Materials and methods

### The dataset

Preprocessed magnetic resource images (MRI) of healthy young adults, comprising structural T1 Weighted Image (T1WI) and diffusion-weighted images, were acquired from the HCP database (Van Essen et al., 2013). The multi-shell diffusion scheme was collected on a 3T scanner with a 64-channel, tight-fitting, brain-array coil. The b-values were 1000, 2000, and 2995 sec/mm^2^. The number of diffusion sampling directions were 90, 90, and 90, respectively. The in-plane resolution was 1.25 mm, and the slice thickness was 1.25 mm (Sotiropoulos et al., 2013).

### GQI tractography

The diffusion data was reconstructed, using GQI tractography, with a diffusion sampling length ratio of 1.25 (F. C. Yeh, Wedeen, & Tseng, 2010). A deterministic, fiber-tracking algorithm was also applied (F.-C. Yeh, Verstynen, Wang, Fernández-Miranda, & Tseng, 2013). Tractography was conducted using the DSI studio software tool (http://dsi-studio.labsolver.org), and tractography seeds were created using the semi-automated FreeSurfer DK cortical atlas (Desikan et al., 2006). The atlas for each subject was imported from the HCP database together with the images. The seed region was created by either the left or the right cerebral white matter (Zajac, Koo, Bauer, Killiany, & Behalf Of The Alzheimer’s Disease Neuroimaging, 2017) with a setting of 10 fixed seeds per voxel (Cheng et al., 2012). The end region was placed on the same side of the seed and was produced by merging the rMFG and the FP into a single region (Fig. 2 A). The normalized quantitative anisotropy threshold was set to 0.075, and the angular threshold was set to 45 degrees. Fiber trajectories were smoothed by averaging the propagation direction with 20% of the previous direction. After confirming the relevant streamline, appropriate indices were obtained using the DSI studio software tool, which included the streamline volume and the diffusion tensor imaging FA mean. A streamline’s volume ratio was added by calculating the streamline volume percentage of the whole white matter volume.

### Spatial location

Constructed streamlines were converted to 3D regions of interest (ROI) using the DSI studio’s tract-to-ROI tool and were registered to an MNI space using the FSL package (FSL, The University of Oxford, Oxford, UK). Then, an overlap image was created using MRIcron software (www.nitrc.org/projects/mricron, University of Nottingham School of Psychology, Nottingham, UK), with all subjects’ SFLT data plotted over a standard MNI template to view common spatial locations and frequencies.

### SFLT subcomponents

The constructed streamline termination points were examined for each subject in the native space using the DK atlas. Accordingly, the SFLT was broken down into subcomponents based on these termination points.

### Spatial relations with adjacent white matter bundles

To study the spatial relationship of the SFLT, a virtual dissection of the brain’s streamlines, generated from an already published HCP-1021 template using DSI studio, was conducted. The HCP-1021 template was averaged from a total of 1021 subjects from the HCP data (F.-C. Yeh et al., 2018). Additionally, to confirm the spatial relation between the SLFT and SFL, as an example, we constructed those tracts in four subjects. The construction methods of the SLF was already described by Wang et. al(Wang et al., 2016).

### The laterality index (LI)

The LI of the SFLT volume ratio and FA mean was calculated using the equation 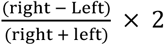. Values between 2 and 0.4 were considered to be right sided, those between −0.4 and −2 were left sided, and values below 0.4 and above −0.4 were undetermined (Catani et al., 2007).

#### Statistical analyses of GQI indices and the LI

Descriptive data is expressed as mean ± standard deviation. A Chi-squared test or Fisher’s exact probability test was applied to assess the correlation of the LI patterns (right, left, and undetermined) of the SFLT volume ratio and the FA mean with the subjects’ gender and handedness groups. Using a two-step cluster analysis method, the SFLT was categorized according to the occurrence of its subcomponents on both sides. Fisher’s exact test assessed the correlation of the SFLT types with the subjects’ gender and handedness groups. Significance in all tests was set at p < 0.05. Statistical analyses were conducted using SPSS (version 25.0) and online statistical software from VassarStats (http://vassarstats.net).

### White-fiber dissection

With the prior knowledge of the GQI tractography results, this study conducted limited brain white-fiber dissections. All brain samples were prepared and fixed according to the method described by Klingler et al. (Klingler and Ludwig 1956). Two right hemispheres from anonymized donors were donated by the Department of Neuroanatomy at Fukushima Medical University, Japan. The study was approved by Fukushima Medical University’s (No. 2137) local review board.

## Abbreviation

AF: Arcuate fasciculus
APF: Anterior projection fibers
BA: Brodmann area
CB: Cingulate bundle
CC: Corpus callosum
cMFG: Caudal middle frontal gyrus
CST: Corticospinal tract
DK: Desikan-Killiany
DLPFC: Dorsolateral prefrontal cortex
DWI: Diffusion weighted image
FA: Fraction anisotropy
FAT: Frontal aslant tract
FLS: Frontal longitudinal system
fMRI: Function magnetic image
FP: Frontal pole
GQI: Generalized q-imaging
HCP: Human connectome project
IFOF: Inferior fronto-occipital fasciculus
LI: Laterality index
MFG: Middle frontal gyrus
MNI: Montreal Neurological Institute
MRI: Magnetic resonance image
PCG: Precentral gyrus
PMd: Dorsal premotor
rMFG: Rostral middle frontal gyrus
ROI: Region of interest
SFG: Superior frontal gyrus
SFLS: Superior frontal longitudinal System
SLF: Superior longitudinal fasciculus
SMA: Supplementary motor area
T1WI: T1 Weighted Image

## Acknowledgments

Brain samples were provided by the Department of Anatomy, Fukushima Medical University. We wish to express our gratitude to the donors, and to Prof. Hiroyuki Yaginuma for his contribution to this research.

## Competing interests

- The authors have no conflicts of interest to disclose.
- This work was partially supported by a Japan Society for the Promotion of Science (JSPS) KAKENHI Grant (number JP18K12110).

The requirement for informed consent was waived due to the nature of the study.

